# The preservation of ancient DNA in archaeological fish bone

**DOI:** 10.1101/2020.04.27.063677

**Authors:** Giada Ferrari, Angelica Cuevas, Agata T. Gondek-Wyrozemska, Rachel Ballantyne, Oliver Kersten, Albína H. Pálsdóttir, Inge van der Jagt, Anne-Karin Hufthammer, Ingrid Ystgaard, Stephen Wickler, Gerald F. Bigelow, Jennifer Harland, Rebecca Nicholson, David Orton, Benoît Clavel, Sanne Boessenkool, James H. Barrett, Bastiaan Star

**Affiliations:** Centre for Ecological and Evolutionary Synthesis, Department of Biosciences, University of Oslo, Blindernveien 31, NO-0371 Oslo, Norway; Norwegian College of Fishery Science, Faculty of Biosciences, Fisheries & Economics, University of Tromsø – The Arctic University of Norway, Tromsø, Norway; McDonald Institute for Archaeological Research, Department of Archaeology, University of Cambridge, Cambridge, CB2 3ER, UK; Cultural Heritage Agency of the Netherlands, Department of Archaeology, Smallepad 5, 3811 MG Amersfoort, The Netherlands; The University Museum, Department of Natural History, University of Bergen, N-5020 Bergen Norway; Department for Historical and Classical Studies, Norwegian University of Science and Technology; The Arctic University Museum of Norway, Tromsø, Norway; Bates College, Pettengill Hall, Lewiston, Maine 04240, USA; Orkney College, University of the Highlands and Islands, East Road, Kirkwall, Orkney, KW15 1LX, UK; Oxford Archaeology, Janus House, Osney Mead, Oxford, OX2 0ES, UK; Department of Archaeology, University of York, York, UK; CNRS, UMR 7209 AASPE, Muséum national d’histoire naturelle (MNHN), CRAVO, Paris, France; Department of Archaeology and Cultural History, NTNU University Museum, 7491 Trondheim, Norway

**Keywords:** Endogenous DNA, bleach, bone element, bone remodeling, petrous bone

## Abstract

The field of ancient DNA is taxonomically dominated by studies focusing on mammals. This taxonomic bias limits our understanding of endogenous DNA preservation for vertebrate taxa with different bone physiology, such as teleost fish. In contrast to most mammalian bone, teleost bone is typically brittle, porous, lightweight and is characterized by a lack of bone remodeling during growth. Using high-throughput shotgun sequencing, we here investigate the preservation of DNA in a range of different bone elements from over 200 archaeological Atlantic cod (*Gadus morhua*) specimens from 38 sites in northern Europe, dating up to 8000 years before present. We observe that the majority of archaeological sites (79%) yield endogenous DNA, with 40% of sites providing samples that contain high levels (> 20%). Library preparation success and levels of endogenous DNA depend mainly on excavation site and pre-extraction laboratory treatment. The use of pre-extraction treatments lowers the rate of library success, although — if successful — the fraction of endogenous DNA can be improved by several orders of magnitude. This trade-off between library preparation success and levels of endogenous DNA allows for alternative extraction strategies depending on the requirements of down-stream analyses and research questions. Finally, we find that — in contrast to mammalian bones — different fish bone elements yield similar levels of endogenous DNA. Our results highlight the overall suitability of archaeological fish bone as a source for ancient DNA and provide novel evidence for a possible role of bone remodeling in the preservation of endogenous DNA across different classes of vertebrates.

## Introduction

Driven by revolutionary advances in laboratory methods, sequencing technologies and downstream analyses, an increasing number of (non-)model species have been investigated using ancient DNA (aDNA). Such studies have addressed a wide range of questions related to archaic human history, animal domestication, or extinct megafauna (e.g., Hofreiter et al., 2015; Ollivier et al., 2018; Palkopoulou et al., 2018; Skoglund & Mathieson, 2018) and have yielded fundamental methodological insights. For instance, a seminal discovery revealed that the petrous bone, i.e., the *pars petrosa* of the temporal bone, which is the hardest and densest bone in mammals (Frisch et al., 1998), has an increased potential of containing high levels of endogenous DNA (Gamba et al., 2014; Pinhasi et al., 2015). Knowing which type of bone element may yield the best results for aDNA research is crucial for a variety of reasons. First, focusing on sampling bone elements with high endogenous DNA greatly improves the economy of high-throughput sequencing studies (Rizzi et al., 2012) and helps avoid costly analyses for samples that are likely suboptimal. Second, sampling for aDNA is most often destructive. Knowing how to select the right elements helps minimize the destruction of unique archaeological materials that represent a finite resource (Pálsdóttir et al., 2019). Third, such knowledge may further aid archaeologists in making informed choices when collecting and preserving zooarchaeological material in the field, maximizing the research potential for a variety of studies. This insight has therefore transformed the field of aDNA, allowing the cost-efficient, genome-wide analysis of hundreds of individual ancient specimens (e.g., Damgaard et al., 2018; Fages et al., 2019; Mathieson et al., 2018; Olalde et al., 2018).

In mammals, low bone density is usually associated with poor DNA preservation (Geigl & Grange, 2018). Archaeological fish bone (Figure 1A) is typically lightweight, porous, brittle and susceptible to taphonomic damage (Szpak, 2011) and such bone could thus be considered a suboptimal source of aDNA from a mammalian preservation perspective. In contrast to mammals, however, fish bone does not serve as a calcium reservoir under normal conditions (Moss, 1961; Witten & Huysseune, 2009). Most higher teleosts, including Atlantic cod (*Gadus morhua*), lack osteocytes (Kranenbarg et al., 2005; Moss, 1961; Shahar & Dean, 2013; Witten & Villwock, 1997). In acellular fish bone, bone remodeling takes place to a lesser extent and through different cellular and physiological processes (Harland & Van Neer, 2018; Kranenbarg et al., 2005; Witten & Villwock, 1997). An absence of bone remodeling may be important for DNA preservation for several reasons. For example, it has been suggested that an absence of cell *lacunae* improves the resistance of acellular fish bone to microbial degradation (Szpak, 2011). Moreover, recent evidence indicates that an absence of bone remodeling may aid DNA preservation in specific mammalian bone elements (Kontopoulos et al., 2019). It is therefore possible that the fundamental differences between mammalian and fish skeletal physiology, and especially the lack of bone remodeling in most fish, affects the aDNA preservation potential of archaeological fish bone.

**Figure 1:**
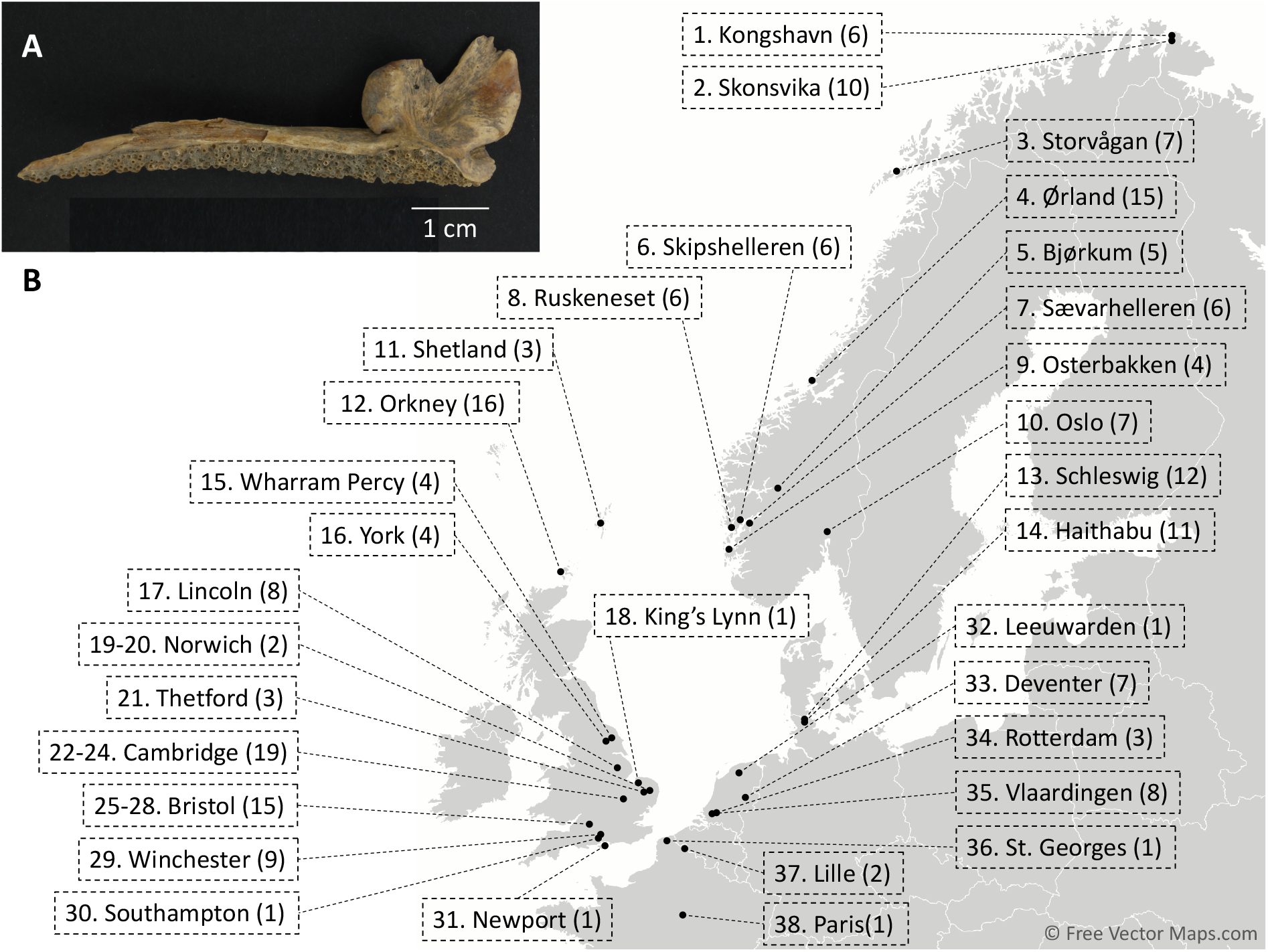
Archaeological Atlantic cod bones. (A) Archaeological Atlantic cod jawbone (premaxilla) from the site of Orkney Quoygrew (1000-1200 CE). (B) Locations of fish bone specimens (*n* = 204) from 38 archaeological sites. Bones from Norwich, Cambridge and Bristol were obtained from distinct, individually numbered archaeological sites (see Table 1).

Interestingly, multiple studies have reported the successful retrieval of aDNA from archaeological fish bone for a variety of species, locations and age (Oosting et al., 2019). aDNA has been consistently amplified from herring (Speller et al., 2012), Pacific salmon (Grier et al., 2013; Johnson et al., 2018; Royle et al., 2018; Speller et al., 2005; Yang et al., 2004), Atlantic cod (Hutchinson et al., 2015; Ólafsdóttir et al., 2014), sturgeon (Ludwig et al., 2009; Nikulina & Schmölcke, 2016; Pagès et al., 2009), Mediterranean trout (Splendiani et al., 2016), Northern pike (Wooller et al., 2015), and other fish taxa (Arndt et al., 2003; Ciesielski & Makowiecki, 2005; Živaljević et al., 2017), in some cases from bones up to 6,000 yBP or older (Johnson et al., 2018; Nikulina & Schmölcke, 2016; Speller et al., 2012; Splendiani et al., 2016; Wooller et al., 2015; Yang et al., 2004). Fish aDNA has also been successfully amplified in metagenomic analyses using bulk bone approaches, even from warm tropical climates (Grealy et al., 2016). Finally, high-throughput sequencing (HTS) approaches have yielded high levels (15-50%) of endogenous DNA from a limited number of sites up to one thousand years old (Boessenkool et al., 2017; Star et al., 2017). Despite the clear potential for aDNA preservation in archaeological fish remains, however, no studies have yet investigated the factors that underlie this preservation and it is unclear if the expectation of intra-skeletal variability in DNA preservation observed for mammals is applicable to other vertebrate taxa such as fish.

Here, we investigate the preservation of aDNA in archaeological Atlantic cod bones (*n* = 204) obtained from 38 excavations in northern Europe, dating from 6500 BCE to c.1650 CE (spanning the Mesolithic to early modern periods, Figure 1B, Tables 1 and S1). We use a HTS approach to investigate whether bone element, archaeological site, DNA extraction method, and/or sequencing library preparation protocol can be used to predict library success (i.e., the successful retrieval and amplification of aDNA) and the relative proportion of endogenous DNA. We interpret our results in light of down-stream analytical requirements and provide practical recommendations in order to maximize throughput and inference of whole genome sequencing (WGS) data from ancient fish bone.

**Table 1:**
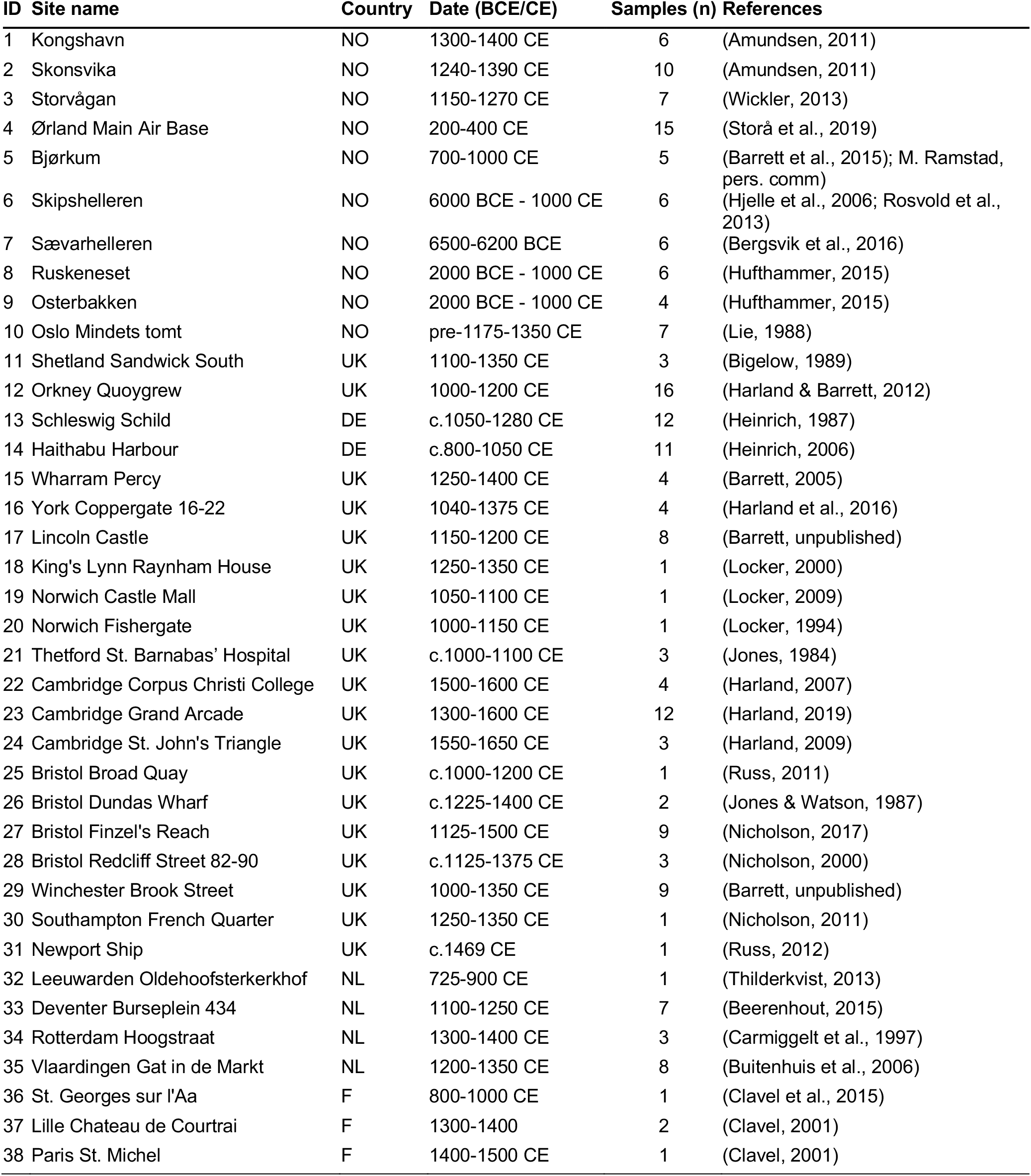
Archaeological sites (*n* = 38) in northwest Europe from which archaeological Atlantic cod bones were obtained. For each site, the country, date and number of bones (*n*) are provided. Dating is based on archaeological context. For locations see also Figure 1B. NO = Norway, UK = United Kingdom, DE = Germany, NL = The Netherlands, F = France.

## Materials and Methods

### Sample processing and DNA extraction

A total of 204 Atlantic cod bones originating from 38 sites (Figure 1B, Tables 1 and S1) were processed following one of three DNA extraction protocols – standard extraction (adapted from Dabney et al., 2013), with the inclusion of a pre-digestion step (DD, Damgaard et al., 2015), or with the addition of a mild bleach treatment and pre-digestion step (BLEDD, Boessenkool et al., 2017). All laboratory protocols were carried out in a dedicated aDNA clean laboratory at the University of Oslo following standard anti-contamination and authentication protocols (e.g., Cooper & Poinar, 2000; Gilbert et al., 2005; Llamas et al., 2017). Bones were UV-treated for 10 minutes per side and pulverized using a stainless-steel mortar (Gondek et al., 2018) or a Retsch MM400 mixer mill. Up to two times 150-200 mg of bone powder was digested for 18-24 hours in 0.5 M EDTA, 0.5 mg/ml proteinase K and 0.5% N-Laurylsarcosine. Digests were combined and DNA was extracted with 9 × volumes of PB buffer (QIAGEN) or a 3:2 mixture of QG buffer (QIAGEN) and isopropanol. MinElute purification was carried out using the QIAvac 24 Plus vacuum manifold system (QIAGEN). Parallel non-template controls were included. A subset of 73 samples was subjected to multiple treatments (Table S1).

### Library preparation, sequencing and read processing

Single- or double-indexed blunt-end sequencing libraries were built from 15-16 μl of DNA extract or non-template extraction blank, following either the single-tube (BEST) protocol (Carøe et al., 2018) with the modifications described in (Mak et al., 2017) or following the Meyer-Kircher protocol (Kircher et al., 2012; Meyer & Kircher, 2010) with the modifications listed in (Schroeder et al., 2015). Blunt-end repair, adapter ligation and set up of indexing PCRs were performed in the aDNA clean laboratory. Library quality and concentration were inspected with a High Sensitivity DNA Assay on the Bioanalyzer 2100 (Agilent) or with a High Sensitivity NGS Fragment Analysis Kit on the Fragment Analyzer™ (Advanced Analytical). Libraries were sequenced on the Illumina HiSeq 2500 or HiSeq 4000 platforms at the Norwegian Sequencing Centre with paired-end 125 bp (HiSeq 2500) or 150 bp (HiSeq 4000) reads and demultiplexed allowing zero mismatches in the index tag. Reads were downsampled (*n* = 100,000) for each library and processed using PALEOMIX v.1.2.13 (Schubert et al., 2014). Paired-end reads were trimmed, filtered, and collapsed with AdapterRemoval v.2.1.7 (Lindgreen, 2012), discarding reads shorter than 25 bp. Collapsed reads were aligned to the Atlantic cod GadMor3 reference genome (Star et al., 2011; Tørresen et al., 2017) with BWA v.0.7.12 (Li & Durbin, 2009), using the aln algorithm with disabled seeding and a minimum quality score of 25. aDNA deamination patterns were assessed with mapDamage v.2.0.6 (Jónsson et al., 2013).

### Statistical analysis

Samples that underwent multiple treatments (*n* = 73, 146 treatments) were used to fit a Generalized Linear Mixed Effect Model (GLM, family = binomial, using sample ID as random effect to account for paired data) to test the effect of DNA extraction protocol, library preparation protocol, site, and bone element on failure or success of library preparation (library outcome ~ extraction protocol + library protocol + site + bone element + (1 | Sample)). Outcome of library preparation was also assessed using all generated libraries, excluding sites with less than three samples (*n* = 191) and controlling for multiple treatments by randomly subsampling one treatment per sample. Subsampling was performed 100 times generating (*i* = 100) resampled datasets. A GLM (library outcome ~ extraction protocol + library protocol + site + bone element) was run on all resampled datasets. A sensitivity analysis was run to evaluate the consistency of the results recording significant factors for each iteration. In order to test the effect of DNA extraction protocol, library preparation protocol, site, and bone element on endogenous DNA content successfully sequenced libraries (defined as libraries that yielded more than 10,000 sequencing reads), excluding sites with less than three successful libraries (*n* = 124 from 19 sites), were used to fit a Generalized Linear Regression (GLR, endogenous DNA fraction ~ extraction protocol + library protocol + site + bone element). Normality of the data for endogenous DNA content was tested by levels in each of the factors using a Shapiro-Wilk Normality Test. For the GLM and GLR described above several models were run discarding factors that did not show significance in more complex models. Akaike (AIC) and Bayesian Information Criterion (BIC) were used to select the best fitting models.

## Results

A total of 277 sequencing libraries were generated from 204 Atlantic cod bones collected at 38 archaeological sites (Figure 1B, Table S1). Of these, 140 libraries from 29 sites had a minimum concentration of 0.1 ng/μl and were sequenced (Figure S1, Figure 3A). All libraries showed patterns of DNA fragmentation, fragment length, and deamination rates that were consistent with those of authentic aDNA (Jónsson et al., 2013; Figures S1 and S2, Table S1). Most samples (*n* = 131) were processed once, but a subset of samples (*n* = 73) was processed using two or more treatment combinations, either using different extraction or library preparation protocols (Figure 3B, Table S1). Bone elements were categorized into three major groups – cranial, postcranial, and pectoral girdle bones (Figure 2A). The representation of these major groups differs across sites (Figure 2B, Table S1), which is driven by the availability of elements at the different locations or by post-excavation sample selection (Box 1).

**Figure 2:**
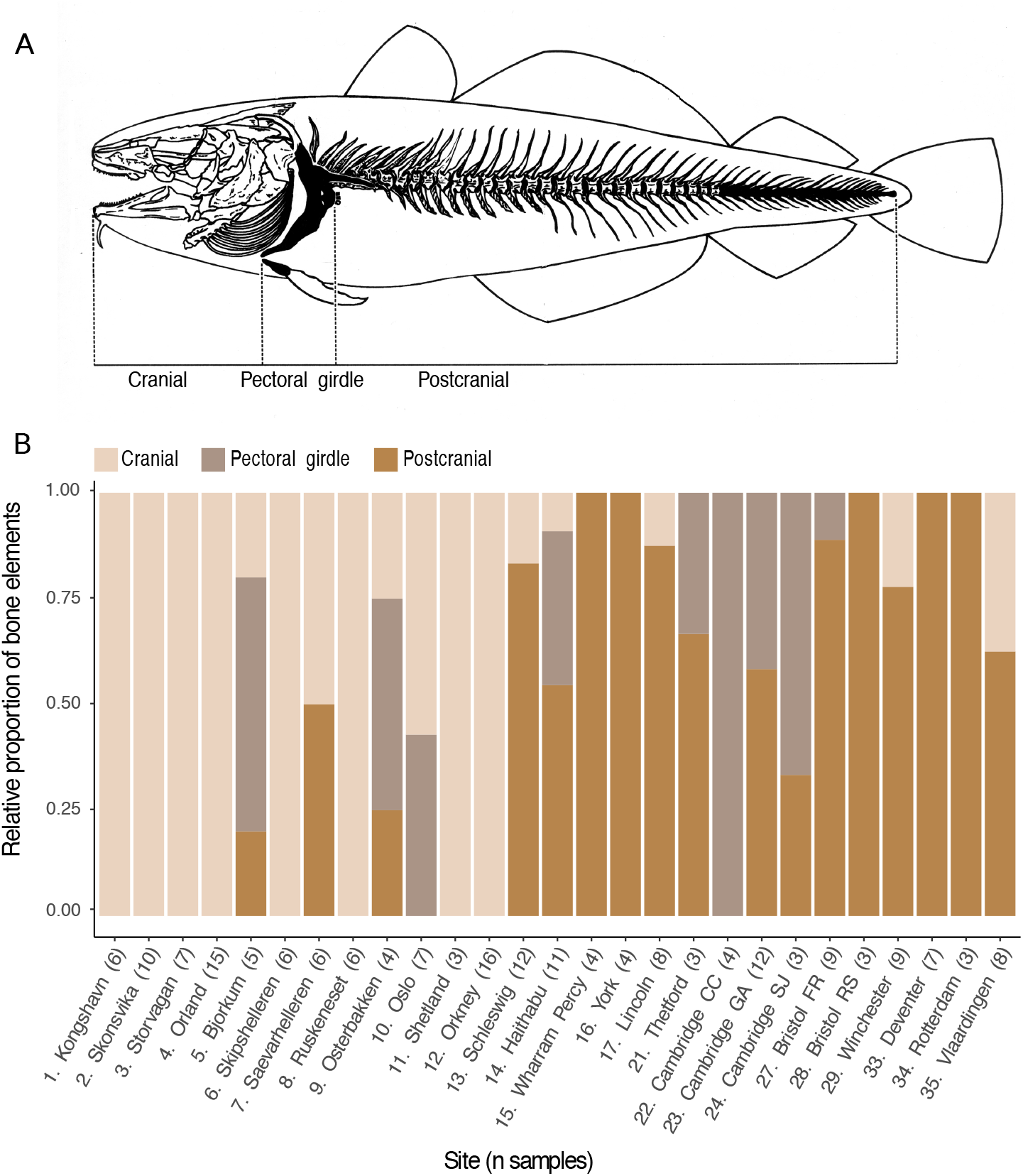
Distribution of Atlantic cod bone elements. (A) Classification of fish bone element groups. Adapted from (Barrett et al., 1999). (B) Distribution of bone elements groups per site. Only sites with three or more samples (n) are shown. Note that the distribution of selected bone elements is not necessarily representative of their relative rate of retrieval at specific sites.

**Figure 3:**
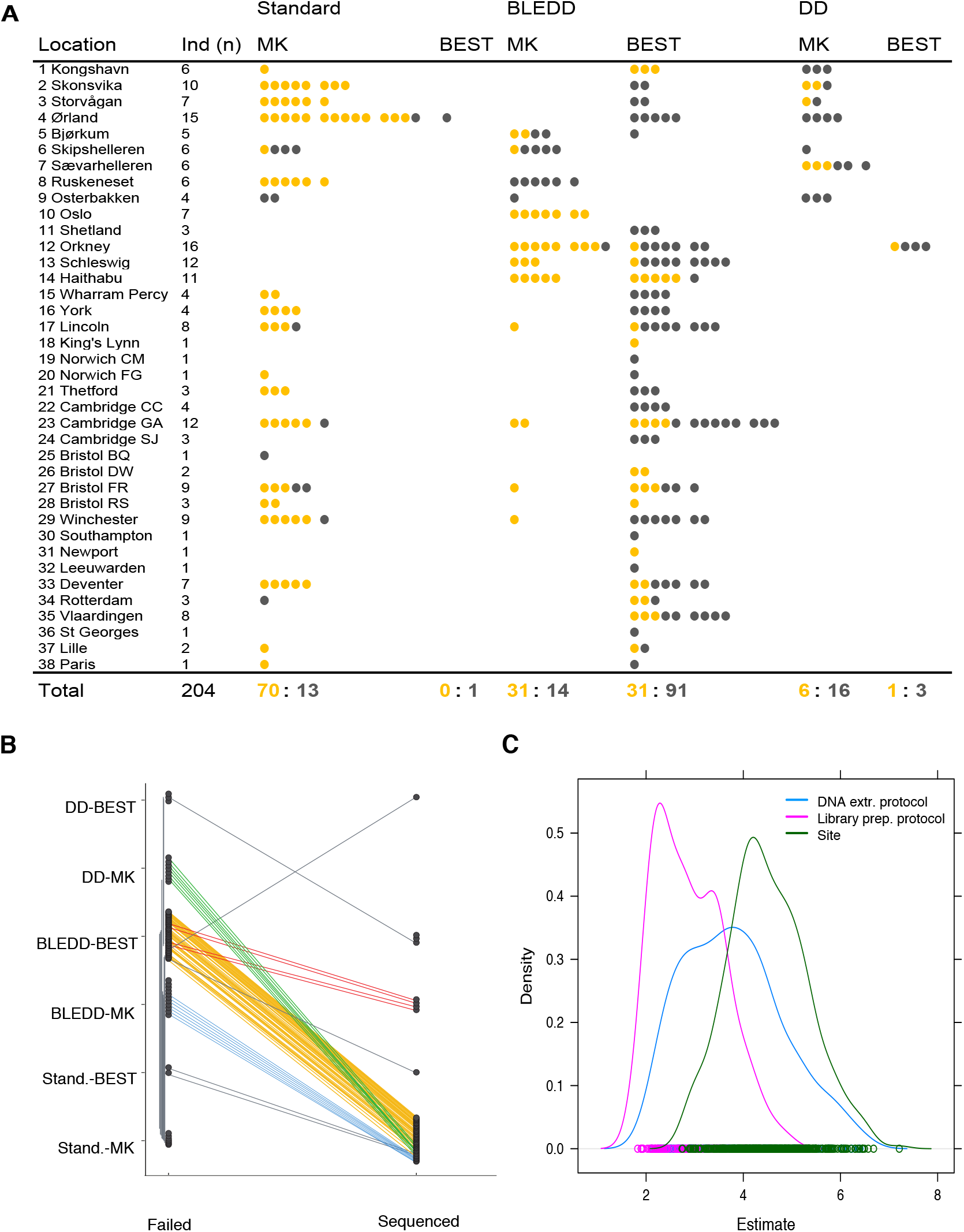
Success rates of high-throughput library preparation from Atlantic cod bones. (A) Schematic of libraries generated for all archaeological sites (yellow, sequenced; gray, processed but not sequenced library) divided into groups according to DNA extraction and library preparation protocols utilized. Note that the number of libraries can be higher than the number of specimens (Ind) due to multiple treatments. DD = double digestion extraction protocol, BLEDD = bleach treatment combined with double digestion extraction protocol, MK = Meyer-Kircher library preparation protocol, BEST = single tube library preparation protocol. (B) Treatment overview and library outcomes (fail or success) for samples processed using multiple treatment combinations (*n* = 73). Different treatment combinations per individual are indicated by connecting lines and colored according to treatment combination. (C) Sensitivity analysis: density distribution of significant factors (site, DNA extraction and library preparation protocols) following iterative (*i* = 100) logistic regression (library outcome ~ extraction protocol + library protocol + site + bone element + (1 | Sample)), using randomly resampled data.

### Box 1

As part of the conservation process prior to long-distance transport, Atlantic cod were typically decapitated (Barrett, 1997) and thus archaeological sites can differ significantly in their bone element distribution (Orton et al., 2014). Specifically, if cod was caught locally, cranial bones may be observed in high abundance, whereas if cod was imported, postcranial bones are likely overrepresented. Bones from the pectoral girdle are anatomically close to the point of decapitation and their presence at import sites may therefore vary (Barrett, 1997; Orton et al., 2014). This variation is clear in the distribution of skeletal elements at different sites in this study (Figure 2B). For example, cod bones found at sites in Norway and Orkney are likely to originate from local catches where cranial bones are abundant. In contrast, sites in England and in the Netherlands are characterized by a lower availability of cranial bones. Moreover, by consistently sampling the same bone element, cranial bones also offer the opportunity to easily avoid resampling the same individual.

Library preparation following the standard extraction protocol generated more successful sequencing libraries (70 out of 84 libraries yielded more than 10,000 sequencing reads) compared to the double digestion (DD) or combined double digestion and bleach (BLEDD) protocols (7 of 26 libraries for DD and 62 of 167 libraries for BLEDD, Figure 3A). The Meyer-Kircher (MK) library preparation protocol yielded a higher success rate (107 out of 150 libraries) than the single tube library protocol (BEST) (32 out of 127 libraries, Figure 3A). A number of library preparations (*n* = 62) for sites with initial high failure rates were repeated using the standard extraction protocol without pre-extraction washes (Figure 3B) resulting in greater success rates. For example, library preparations after DNA extraction using the DD (*n* = 4) or BLEDD (*n* = 5) protocols for samples from the site of Ørland Main Air Base (site 4) failed, while library preparation following the standard DNA extraction protocol was more successful (14 out of 17 libraries, Table S1).

To statistically infer the most important factors explaining library success we applied two models. First, we focused on the samples that were processed with multiple treatments (*n* = 73, Figure 3B). Second, we incorporated all samples generated from sites with more than three samples (*n* = 191 samples from 27 sites), correcting for multiple treatments by randomly downsampling a single treatment per sample iteratively (*i* = 100, Figure 3C). The GLM focusing on the samples with multiple treatments (library outcome ~ extraction protocol + library protocol + site + bone element + (1 | Sample)) shows that the outcome is significantly dependent on DNA extraction and library preparation protocols (Table S2). The sensitivity analysis with the 100 iterations of GLMs incorporating all samples (library outcome ~ extraction protocol + library protocol + site + bone element) shows similar results, with site and DNA extraction protocol as the most prevalent significant factors (presenting mean estimates across iterations of 4.50 and 3.79 respectively, Figure 3C), followed by library preparation protocol (mean estimate = 2.90). Bone element has no significant effect on library preparation outcome and after excluding it from the model the latter shows a better fit to the data (Table S3).

We further assessed whether the same factors affect levels of endogenous DNA for samples (*n* = 124) from 19 locations for which three or more specimens were successfully sequenced by fitting a GLR (endogenous DNA fraction ~ extraction protocol + library protocol + site + bone element). Significantly higher endogenous DNA contents are observed in samples that underwent the DD or BLEDD pre-treatments, compared to a standard DNA extraction (Figure 4A, Table S4). Given that a number of samples for which DD or BLEDD extraction failed (*n* = 62) were re-extracted using the standard protocol (figure 3B), such samples may *a priori* be suspected to have relatively poor DNA preservation. In contrast, library preparation protocol had no significant effect on endogenous DNA content (Table S4). Although postcranial bones tend to have lower levels of endogenous DNA, these differences are not significant, and especially bones from the cranial and pectoral girdle yield comparable levels of endogenous DNA, independent of DNA extraction protocol (Figure 4B, Table S4). Finally, we observe significant differences in endogenous DNA between sites (Figure 4C, Table S4) with 8 out of 19 sites yielding samples with high levels (> 20%) of endogenous DNA, which includes the oldest excavation (Sævarhelleren, site 7, dated to ca. 6500-6200 BCE). When excluding the non-significant factors from the GLR (bone element and library preparation protocol, Table S5), DNA extraction protocol and site remain significant. The most complex model including all factors shows the best fit to the data (Table S5).

**Figure 4:**
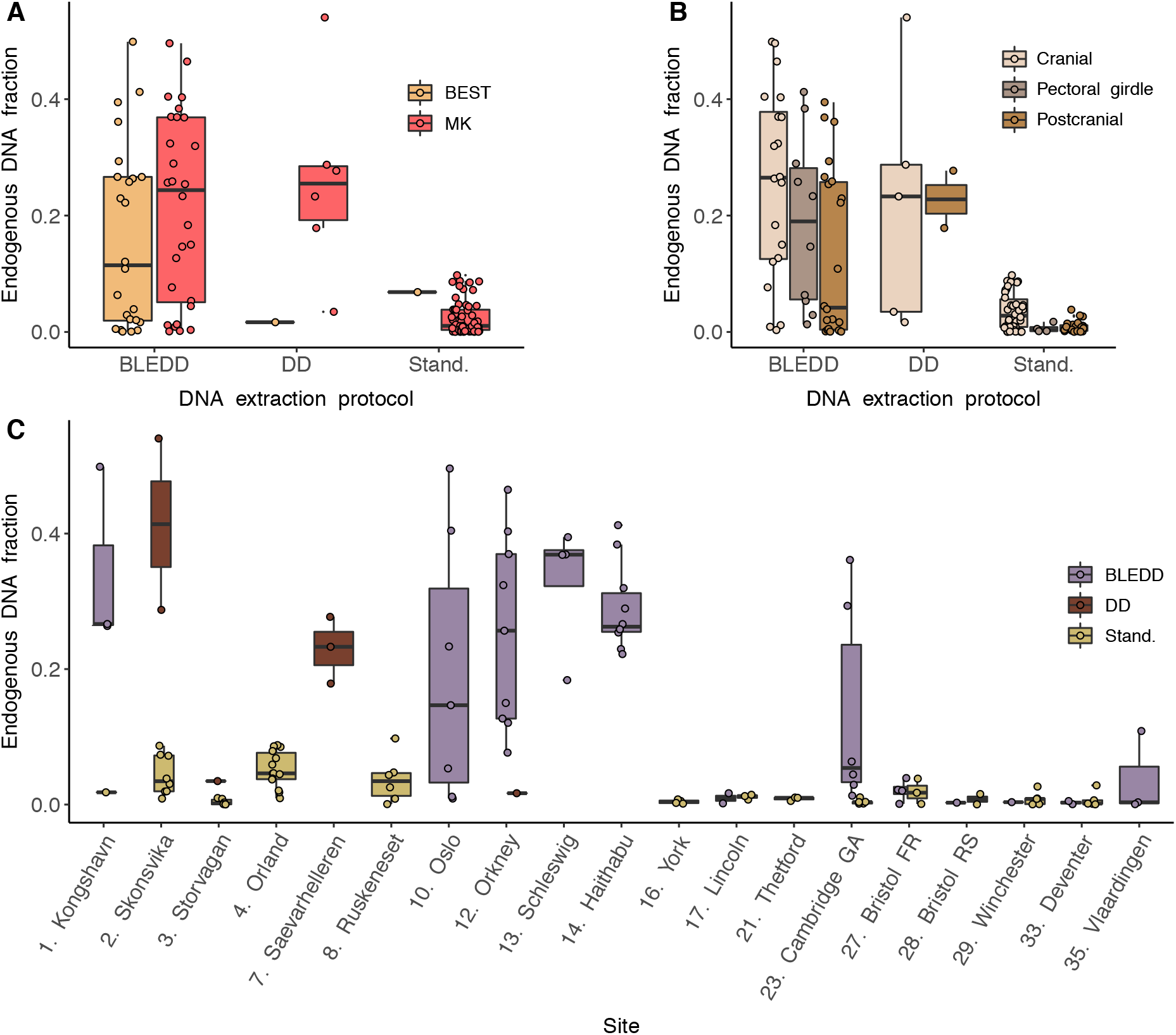
Endogenous DNA fraction. (A) Endogenous DNA per extraction and library preparation protocols. Double digestion and mild bleach wash pre-treatments result in higher endogenous DNA, independently from library preparation protocols. (B) Endogenous DNA per skeletal element. No significant differences in endogenous DNA content can be observed between cranial, postcranial and pectoral girdle bones. (C) Endogenous DNA per site. Significant differences in DNA preservation can be observed between sites. DD = double digestion extraction protocol, BLEDD = bleach treatment and double digestion extraction protocol, MK = Meyer-Kircher library preparation protocol, BEST = single tube library preparation protocol. Only sites for which three or more libraries were successfully sequenced are plotted.

## Discussion

Here, we present the largest study on DNA preservation in ancient fish bones to date, assessing the effects of bone element, archaeological site, DNA extraction and sequencing library preparation protocols on library success and levels of endogenous DNA. We obtain several conclusions.

First, although we did not exhaustively sample all different elements possible, our findings imply that most fish bone elements of sufficient size may be suitable for high-throughput shotgun aDNA analyses. We observe no significant differences in either library preparation success or endogenous DNA levels amongst different bone element groups (i.e., cranial, pectoral girdle or postcranial) in archaeological fish bones. This observation is strikingly different from ancient DNA results obtained from mammalian bones, where high endogenous DNA preservation is localized, either in the petrous bone (Gamba et al., 2014; Pinhasi et al., 2015) or in the dense, recently deposited circumferential lamellae of long bones (Alberti et al., 2018). Moreover, our findings support a recent alternative hypothesis for DNA preservation in vertebrate bone. The localized DNA preservation in mammals has usually been explained by the observed high density of bones or bone regions (Bollongino et al., 2008; Geigl & Grange, 2018; Kendall et al., 2018; Alberti et al., 2018) that may be more resistant to exogenous microbial colonization or taphonomic degradation (Campos et al., 2012; Gamba et al., 2014). Recently, however, it has been suggested that it is the absence of bone remodeling in these bones that helps promote DNA preservation (Kontopoulos et al., 2019; Sirak et al., 2020), following observations that the petrous bone (Kontopoulos et al., 2019), the auditory ossicle (Sirak et al., 2020) and the circumferential lamellae of long bones (Treuting et al., 2017) experience little or no bone remodeling. This absence is analogous to the lack of bone remodeling during growth in acellular fish bone (Kranenbarg et al., 2005; Witten & Villwock, 1997). Here, we provide novel comparative data supporting a hypothesis that distinct bone developmental characteristics, specifically bone remodeling (e.g., Kontopoulos et al., 2019; Sirak et al., 2020), may contribute to increased levels of endogenous DNA preservation in a wide range of vertebrates.

Second, depending on the down-stream computational requirements, sample sizes consisting of poor-quality DNA specimens can be increased in an economical way by avoiding pre-extraction digestion or bleach wash treatments. We observed a distinct trade-off between levels of endogenous DNA and library success when using bleach wash and pre-digestion treatments. As previously reported, bleach wash and pre-digestion treatments increase levels of endogenous DNA (e.g., (Boessenkool et al., 2017; Damgaard et al., 2015; Korlević et al., 2015), yet this increase is coupled to higher failure rates during library creation. Presumably, when samples have relatively poor DNA preservation, this DNA can be lost using pre-extracting wash steps, resulting in the failure of a sample that could otherwise yield a sequencing library with low levels of endogenous DNA. This results in a trade-off where the number of investigated individuals can be maximized at the cost of sequencing depth or vice-versa. Such a trade-off can be exploited in situations where low sequencing coverage data can yield meaningful archaeological or biological information. For instance, genetic sex can be easily obtained for mammals using low numbers (e.g., < 10,000) of sequencing reads, even in samples with low levels (< 0.5%) of endogenous DNA (e.g., Barrett et al., 2020; Nistelberger et al., 2019; Pečnerová et al., 2017). Analogously, in Atlantic cod there are several large, ecologically important chromosomal inversions (e.g., Berg et al., 2016; Berg et al., 2017) that can easily be determined using low coverage sequencing data (Star et al., 2017).

Third, we recommend utilizing library protocols with intermediate purification procedures when targeting archaeological fish bones in order to maximize the potential of successful library creation. It is advantageous to minimize hands-on-time and laboratory costs while simultaneously increasing sample throughput. For this reason, we implemented the single-tube (BEST) protocol (Carøe et al., 2018; Mak et al., 2017), which is an economically efficient protocol with a reduced number of purification steps compared to the Meyer-Kircher protocol (Kircher et al., 2012; Meyer & Kircher, 2010). Both protocols yield similar levels of endogenous DNA and can therefore be used to retrieve high-quality aDNA libraries from archaeological fish bone. Nonetheless, we did observe significantly increased library amplification failure rates when following BEST (Carøe et al., 2018; Mak et al., 2017), which reduces the efficiency of this method in overall sample throughput. During the BEST protocol, multiple enzymatic reactions occur successively in the same tube, and we suspect it is possible that this protocol is more sensitive to contaminants than protocols with intermediate purification steps such as the Meyer-Kircher protocol. It is further possible that archaeological fish bone contains more such contaminants than mammalian bone. Fish bone may therefore be less suited for single-tube library preparation protocols than mammalian bone, which can be more efficiently cleaned.

Finally, we conclude that a wide range of preservation and excavation conditions can yield high endogenous aDNA preservation in archaeological fish bone. We observe site-specific differences in aDNA preservation, with some sites yielding consistently high rates of library success and levels of endogenous DNA whereas others do not. These site-dependent results make it difficult to predict specific factors underlying sufficient aDNA preservation, as samples from each site are associated with a wide range of different, potentially unknown, *pre*- and *post*-excavation taphonomic processes. However, our results confirm that cave sites typically offer ideal conditions for DNA preservation (Bollongino et al., 2008; Hardy et al., 1995), thanks to stable low temperatures and lack of precipitation (Hedges & Millard, 1995). Here, we report the oldest WGS results for archaeological fish bone from the cave site of Sævarhelleren (site 7, Bergsvik et al., 2016), which is one of the sites with better DNA preservation despite being up to 8500 years old. In addition to this, we have obtained excellent DNA of bones obtained from dry shell middens (e.g., Orkney Quoygrew, site 12, Harland & Barrett, 2012), as well as bones from waterlogged sediments that were excavated decades ago (e.g., Haithabu Harbour, site 14, Heinrich, 2006).

Only a limited number of fish aDNA studies have been published, despite their environmental and economic importance, and the abundance of archaeological fish bone (Barrett, 2019; Oosting et al., 2019). Especially whole genome HTS approaches (e.g., Star et al., 2017) focusing on fish remains are rare. Here we show that, despite high variability in DNA preservation across archaeological sites, high endogenous aDNA can consistently be recovered from archaeological fish bone. Overall, we obtained successful sequencing libraries from 50% of all fish bone samples analyzed, retrieving samples with more than 20% endogenous DNA from 40% of sites. Our results provide insights for the study design and laboratory processing of archaeological fish bone remains, highlight the suitability of this material as a source for aDNA, and provide novel evidence for the role of bone remodeling in the preservation of DNA in vertebrate bone.

## Data availability

All ancient read data are available at the European Nucleotide Archive, www.ebi.ac.uk/ena, (accession no. PRJEB37681).

## Acknowledgments

We thank C. Amundsen for providing the Kongshavn and Skonsvika bone samples, M. Skage, S. Kollias and A. Tooming-Klunderud at the Norwegian Sequencing Centre for sequencing and processing of samples, and K. Dean for comments on the statistical analyses. This work was supported by Research Council of Norway Project “Catching the Past: Discovering the legacy of Atlantic cod exploitation using ancient DNA” (262777) and Leverhulme Trust Project “Northern Journeys: Reimagining the Medieval Revolution and its Aftermath” (MRF-2013-065). Analyses were performed on the Abel Cluster at the University of Oslo and the Norwegian metacentre for High Performance Computing (NOTUR), operated by the Department for Research Computing at the University of Oslo IT-department (USIT).

## Author contributions

B.S., G.F. and J.H.B designed research; O.K., A.H.P., A.T.G. and G.F. carried out laboratory work. A.C. and G.F. analyzed data. A.K.H., I.Y., I.J., S.W., G.F.B., J.H., R.N., D.O., B.C., R.B. and J.H.B. provided samples and archaeological context information. R.B. selected UK specimens. B.S. and G.F. wrote the manuscript with input from all authors. B.S., J.H.B. and S.B. provided funding and consumables.

